# Myofibril and mitochondria morphogenesis are coordinated by a mechanical feedback mechanism in muscle

**DOI:** 10.1101/2020.07.18.209957

**Authors:** Jerome Avellaneda, Clement Rodier, Fabrice Daian, Thomas Rival, Nuno Miguel Luis, Frank Schnorrer

**Affiliations:** Aix Marseille University, CNRS, IBDM, Turing Center for Living Systems, 13288 Marseille, France

**Keywords:** muscle, *Drosophila*, mitochondria, mitochondrial fusion, sarcomere, fate determination, biomechanics

## Abstract

Complex animals build specialised muscle to match specific biomechanical and energetic needs. Hence, composition and architecture of sarcomeres as well as mitochondria are muscle type specific. However, mechanisms coordinating mitochondria with sarcomere morphogenesis are elusive. Here we use *Drosophila* muscles to demonstrate that myofibril and mitochondria morphogenesis are intimately linked. In flight muscles, the muscle selector *spalt* instructs mitochondria to intercalate between myofibrils, which in turn mechanically constrain mitochondria into elongated shapes. Conversely in cross-striated muscles, mitochondria networks surround myofibril bundles, contacting myofibrils only with thin extensions. To investigate the mechanism causing these differences, we manipulated mitochondrial dynamics and found that increased mitochondrial fusion during myofibril assembly prevents mitochondrial intercalation in flight muscles. Strikingly, this coincides with the expression of cross-striated muscle specific sarcomeric proteins. Consequently, flight muscle myofibrils convert towards a partially cross-striated architecture. Together, these data suggest a biomechanical feedback mechanism downstream of *spalt* synchronizing mitochondria with myofibril morphogenesis.

## Introduction

Muscles power all voluntary animal movements. These movements are produced by arrays of myosin motors that are assembled together with titin and actin filaments into elaborate contractile machines called sarcomeres^1,2^. Hundreds of sarcomeres are connected into long chains called myofibrils that span the entire muscle fiber and thus mechanically connect two skeletal elements^3^. During muscle contraction each myosin motor head consumes one molecule of ATP per crossbridge cycle to move myosin about 10 nm relative to actin and produce a few piconewton of force^4^. Thus, sustained muscle contraction requires large amounts of ATP.

As ATP is most effectively produced by oxidative phosphorylation in mitochondria, muscles generally contain large amounts of mitochondria. However, mitochondrial content varies to a large extent between different muscle types and across species^5^, suggesting that mitochondria biogenesis is adjusted to match the energetic requirements of muscle fiber types. A striking example are slow oxidative muscle fibers of mammals that are enduring muscles and thus strongly depend on high ATP levels. These fibers contain larger amounts of mitochondria compared to fast glycolytic fibers^6^. However, not only total mitochondrial content but also mitochondrial morphology is fiber-type dependent with more elongated mitochondria present in slow fiber types^6^. This suggests that mitochondria biogenesis is intimately linked to muscle fiber type-specific physiology. However, the molecular mechanisms of this coordination are unclear.

Recent advances in high resolution imaging revealed that mitochondrial morphologies in individual muscle fibers are not homogeneous. Mitochondria closer to the plasma membrane are generally more globular, whereas mitochondria in proximity to myofibrils are part of more complex networks^7^. Parts of the mitochondrial network contact the sarcomeric I-bands, other parts run in parallel to the fiber axis, in close proximity to the myofibrils^8^. Strikingly, the organisation of mitochondrial networks also depends on the muscle fiber type: oxidative fibers contain more mitochondria preferentially oriented in proximity to and in the direction of myofibrils, a phenomenon even more prominent in the heart, a muscle that strictly depends on ATP production by oxidative phosphorylation^9^. Hence, ATP production is located close to the ATP consuming contractile motors. However, little is known about the mechanisms of how myofibril and mitochondria development are coordinated to match the energetic requirements with the contractile properties of muscle fibers.

To investigate the interplay between myofibrils and mitochondria we turned to *Drosophila* and compared two different muscle types, indirect flight muscles and leg muscles. Indirect flight muscles of insects are specialised to combine high power output with endurance and thus use oxidative metabolism. *Drosophila* flight muscles oscillate at 200 Hz and produce up to 80 Watt power per kg of muscle mass during long flight periods^10-12^. This is achieved by a fibrillar morphology of their contractile myofibrils enabling a stretch-activation mechanism to trigger the fast oscillations^13^. Hence, the ATP demand of these muscle fibers during flight is extremely high. With its strict aerobic metabolism and its stretch activation mechanism requiring high mechanical tension, insect flight muscles biomechanically and energetically resemble the mammalian heart muscle^14,15^.

In contrast, the other adult *Drosophila* body muscles found in legs or abdomen show a regular cross-striated myofibril morphology resembling mammalian skeletal muscle fibers and use a normal synchronous contraction mechanism^16,17^. Thus, their energy requirements are strikingly different from flight muscles.

Here, we compared myofibril and mitochondria morphologies between flight and leg muscles of *Drosophila* and found that flight muscle mitochondria are mechanically squeezed against myofibrils maximising their contact areas and isolating neighbouring myofibrils. We discovered that mitochondrial intercalation between myofibrils coincides with myofibril assembly. Strikingly, if intercalation is prevented by increased mitochondrial fusion fibrillar flight muscles express sarcomeric proteins specific to the cross-striated leg muscle type resulting in a partial conversion to cross-striated fiber morphology. This suggests a mechanical interplay between mitochondria dynamics and myofibril development, which triggers a feedback mechanism coordinating mitochondria with myofibril morphogenesis.

## Results

### Muscle type specific mitochondria morphology is instructed by Spalt

In order to examine the regulation of mitochondria biogenesis and myofibril morphogenesis in different fiber types, we chose *Drosophila* adult indirect flight muscles and leg muscles as models. We visualised myofibril morphology with phalloidin and mitochondria morphology by expressing GFP fused to a mitochondrial matrix targeting signal (mito-GFP) with *Mef2-*GAL4. Flight muscles show the expected fibrillar myofibril morphology (Fig. 1a,b’)^17^. Flight muscle mitochondria are densely packed around the individual myofibrils, adopting an elongated shape along the myofibril axis, consequently physically isolating neighbouring myofibrils (Fig. 1b,b’’).

**Fig. 1.**
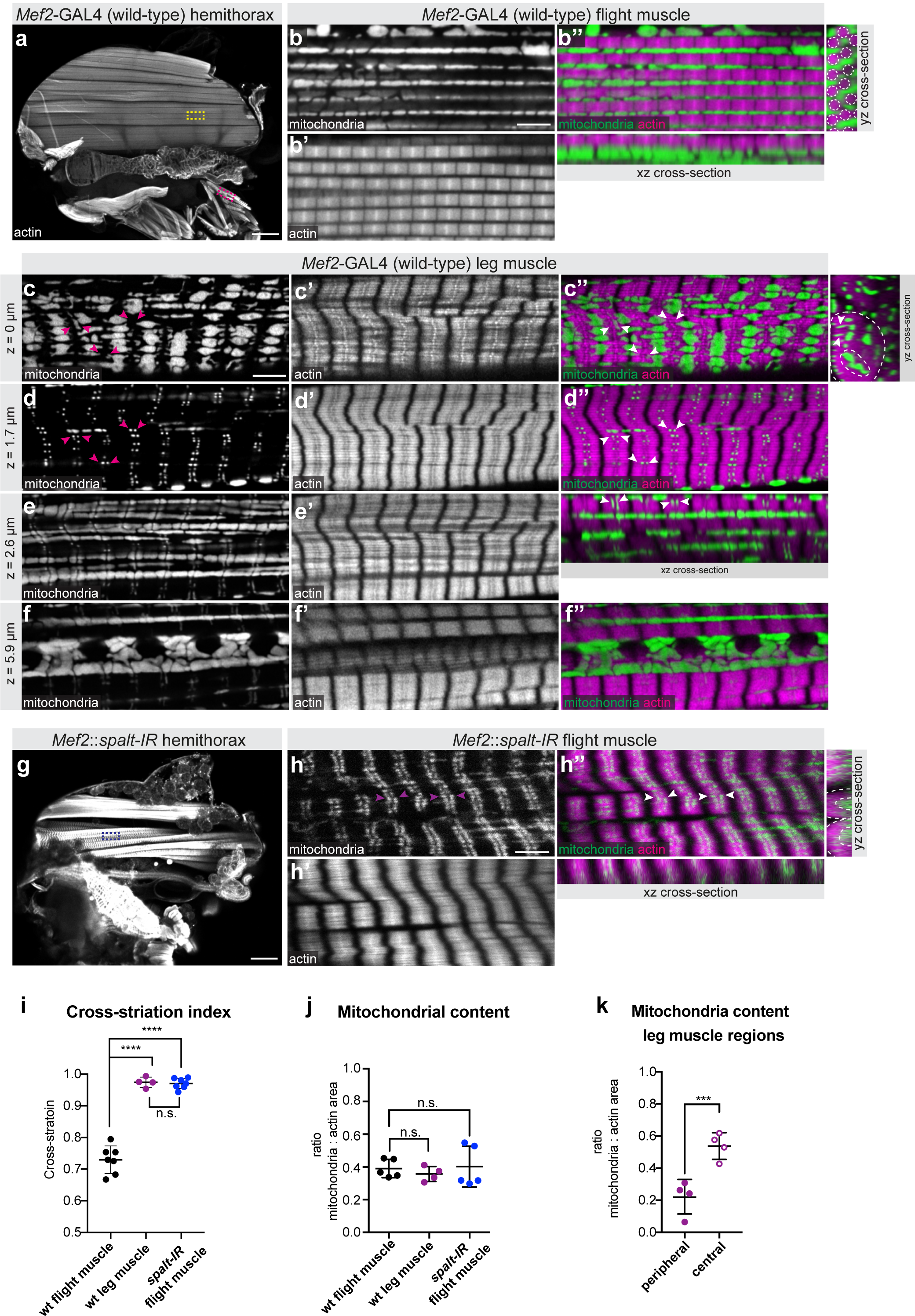
*spalt* regulates muscle type-specific mitochondria morphogenesis. (**a-f**) *Mef2-*GAL4 (wild-type) hemi-thorax (a), flight muscle (b) and leg muscle (c-f) stained with phalloidin to visualise actin (magenta) and expressing mito-GFP to visualise the mitochondrial matrix (green). Yellow and magenta boxes in (a) indicate representative regions of flight and leg muscles magnified in (b-f). Single confocal plane as well as and xz yz cross-sections are shown (b’’). Note the individualised myofibrils (dotted circles) surrounded by densely packed mitochondria. (c-f) Leg muscle top (c), middle (d,e) and central slice (f) a showing the tubular fiber morphology (yz cross section), cross-striated myofibrils and complex mitochondrial shapes filling the surface and the center of the myofiber and contacting the sarcomeric I-band with thin extensions (magenta and white arrow heads). (**g-h**) *Mef2::spalt-IR* hemithorax (g) and flight muscle (h) display tubular fiber morphology (h’’ yz cross-section), cross-striated myofibrils and centrally located mitochondria with thin extension towards the I-band (arrow heads). (**i-k**) Quantification of the lateral fibrillar alignment called cross-striation index in muscle (i, see Supplementary Fig. 1), the relative mitochondria content (j, relative to myofibril content) and the mitochondria content in leg muscle regions (k). Dotted lines in cross sections in (c’’) and (h’’) represent the regions measured. Note that higher mitochondria density in the center of leg muscles. In all plots, individual circles represent individual animals, for each a minimum of 5 measurements was done, and mean +/- standard-deviation (SD) is indicated. Significance from unpaired t-tests is denoted as p-values p ≤ 0,001 (***) or ≤ 0,0001 (****). (n.s.) non-significant. Scale bars are 100 µm (a,g) and 5 µm (b,c,h).

In contrast, leg muscles have cross-striated myofibrils, which align to form a tube, whose center is devoid of myofibrils (Fig. 1a,c’-f’,i)^17^. Interestingly, leg muscle mitochondria do not intercalate within the cross-striated myofibrils, but are present both peripherally (Fig. 1c) and centrally, where they are strongly concentrated (Fig. 1f,k). They appear to be largely excluded from the area occupied by the cross-striated myofibrils, with only small mitochondrial extensions contacting the sarcomeric I-band (Fig. 1d,e). Such a specific mitochondrial-myofibril contact area is not found in the fibrillar flight muscles, however, the overall mitochondrial content, when normalised to the actin content of flight and leg muscles is comparable (Fig. 1j).

The formation of fibrillar flight muscle requires the zinc-finger transcription factor Spalt (Spalt major, Salm)^17^. Interestingly, we found that knock-down of *spalt* in flight muscles during development using *Mef2-*GAL4 not only transforms myofibrils into a cross-striated tubular morphology (Fig. 1g-i, Supplementary Fig. 1) but also converts the simpler mitochondrial morphology of flight muscles into a leg-specific type with centrally concentrated mitochondria, which contact the sarcomeric I-bands with thin extensions (Fig. 1h). Taken together, these data show that the physiologically and mechanically distinct muscle fiber types of adult flies display strikingly different mitochondrial morphologies. In flight muscles, mitochondria and myofibrils morphologies are both instructed by the transcriptional regulator Spalt.

### Flight muscle mitochondria elongate in proximity to myofibrils

In order to examine in more detail mitochondria morphology in relation to myofibril structure, we developed a method to better quantify mitochondria morphologies in the different muscle types. To be able to delineate mitochondrial shape in an automated way in flight muscles, we established a live dissection method avoiding fixation. Additionally, we generated a new marker line labelling the mitochondrial outer membrane by fusing GFP to the mitochondrial outer membrane localisation signal of Tom20, here named MOM-GFP (see Methods). When expressed in flight muscles with *Mef2-*GAL4, MOM-GFP delineates the mitochondrial outer membrane (Fig. 2a), which enabled us to segment and reconstruct individual mitochondria in three dimensions using a deep learning network (Fig. 2b,c, Supplementary Fig. 2, Supplementary Movie 1). These data show that the average volume of flight muscle mitochondria is about 3-4 µm^3^ (Fig. 2d,f). Most mitochondria adopt a simple elongated ellipsoid-like shape with the long axis of the ellipsoid oriented in the direction of the myofibrils (Fig. 2e-g). Specific extensions towards the myofibrils are absent, instead very large contact areas between myofibrils and mitochondria are likely present in flight muscle.

**Fig. 2.**
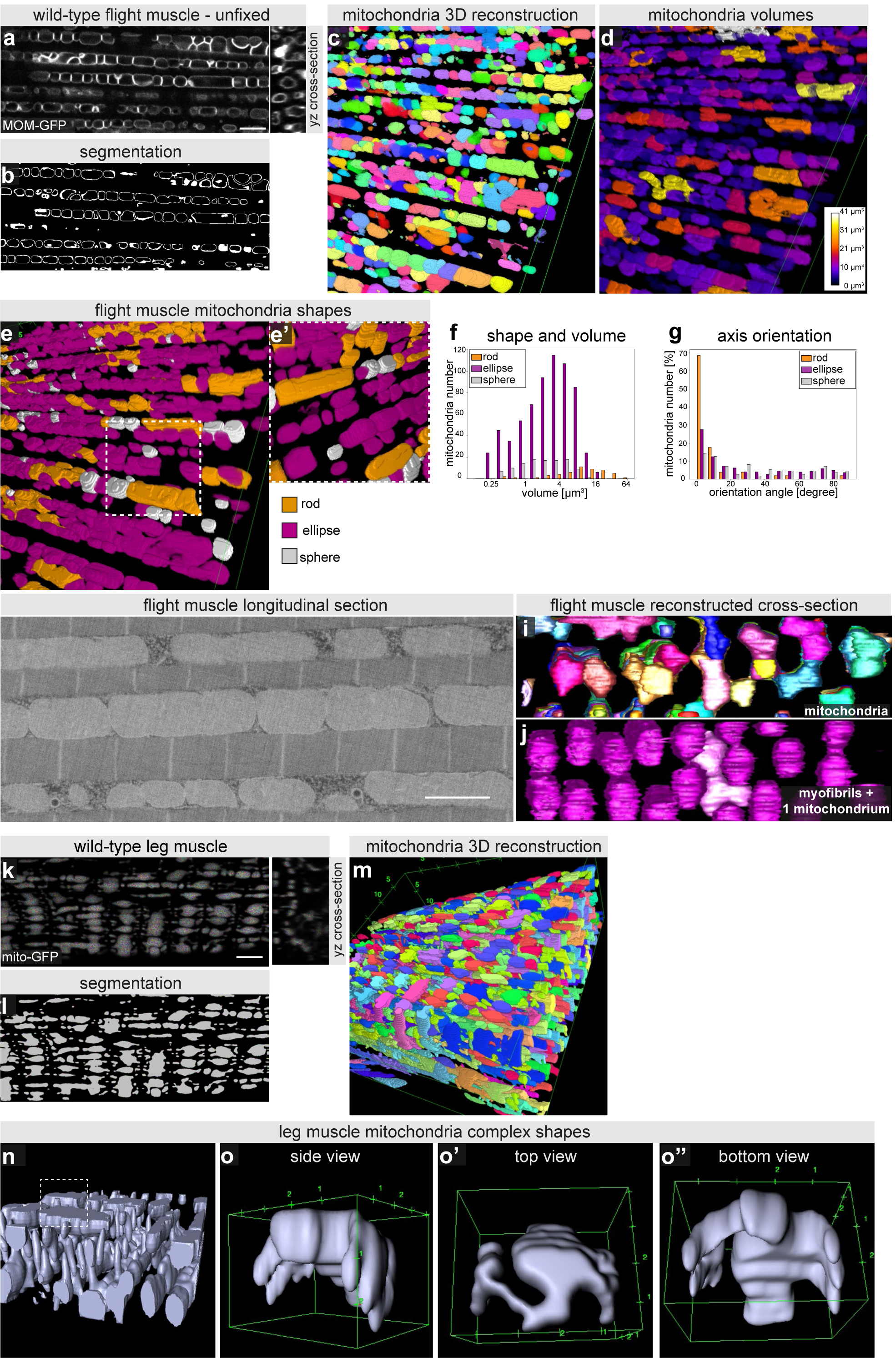
Quantification of mitochondrial morphology in muscle types. (**a-g**) Highly resolved confocal sections of unfixed alive flight muscle mitochondria labelled with MOM-GFP expressed with *Mef2-*GAL4 (a) to segment the mitochondria outlines using machine learning (b, see Supplementary Fig. 2). 3D segmentation of individual flight muscle mitochondria using Fiji (c), each mitochondrion was assigned a random colour (see Supplementary Movie 1). Classification of individual mitochondria based on their volume (d) and shape classifiers (e); close up (e’), see Methods for the classification parameters. Total mitochondria number and their volumes in a 67.5 µm x 67.5 µm x 6.7 µm volume (f). Note the preferred orientation of the long mitochondrial axis with the axis of the myofibrils (g). (**h-j**) Serial block face electron microscopy of adult flight muscles, showing a longitudinal view (h). Note the intimated contact of mitochondria and myofibrils. Cross-section view of a 3D-reconstruction of individual mitochondria shown in different colours (i) and of the myofibrils in magenta with one mitochondrion in light pink (j, Supplementary Movie 2). Note the mitochondrial indentations caused by pushing myofibrils. (**k-m**) Fixed leg muscle mitochondria labelled with mitochondrial matrix GFP (mito-GFP) expressed with *Mef2-*GAL4 (Supplementary Movie 3). A representative peripheral (top) section of the z-stack (used in Fig. 1c) and a yz-cross-section orthogonal view are show (k). Interactive Watershed using Fiji allowed segmentation (i) and 3D reconstruction of individual mitochondria (j, Supplementary Movie 4). (**o-p**) 3D reconstruction of a cropped leg muscle volume shows complex mitochondria morphologies with thin extensions from surface and central mitochondria into the myofibril layer (o, Supplementary Movie 5). Side-, top- and bottom-views of an individual mitochondrion displaying its complex morphology (p-p’’, Supplementary Movie 6). Scale bars are 5 µm in a,k and 2 µm in h.

To resolve these contact areas in more detail we applied serial block-face electron microscopy and indeed could verify the intimate contacts with virtually no detectable space between mitochondria and myofibrils (Fig. 2h, Supplementary Movie 2). By reconstructing myofibrils and mitochondria in three dimensions we found that the majority of elongated mitochondria are squeezed against individual myofibrils resulting in round indentations in the mitochondria that cover about half of a myofibril circumference (Fig. 2j, Supplementary Movie 2). Mitochondria do not form networks but are rather individualised with an average volume of an individual mitochondrion of 3.9 µm^3^, which is in good accordance with our light microscopy quantifications. These data demonstrate that myofibril and mitochondria morphologies are intimately linked in flight muscles and thus suggest that myofibril development is highly coordinated with mitochondria morphogenesis.

### Leg muscle mitochondria acquire complex shapes

To segment the complex shapes of leg muscle mitochondria we used the mitochondrial matrix mito-GFP marker expressed with *Mef2-*GAL4 (Fig. 2j,k, Supplementary Movie 3). This allowed us to attempt a 3D reconstruction of leg muscle mitochondria (Fig. 2l,m, Supplementary Movie 4). However, the success of the automated segmentation of individual mitochondria was limited by their complex shapes and thin extensions. Manual reconstruction displayed these complex shapes with elongated structures extending in 3D, and particularly prominent extensions towards the sarcomeric I-bands (Fig. 2n,o, Supplementary Movies 5,6). Likely, our manual segmentation underestimates the connectivity between these complex mitochondria, as the detection of thin connections by light microscopy is limited. Thus, leg muscle mitochondria organise into rather complex networks above and below the aligned cross-striated myofibrils, and hence are strikingly different from flight muscle mitochondria.

### Flight muscle mitochondria morphology is ruled by mechanical pressure

We found that flight muscle mitochondria are in intimate contact with myofibrils and acquire an elongated shape. We have shown in the past that myofibrils are under significant mechanical tension during development and that this tension is required to build linear myofibrils^18,19^. Hence, we hypothesized that myofibril tension creates a pushing force against mitochondria that constrain them into the observed ellipsoid shape. To test this hypothesis, we applied our live dissection protocol of flight muscles combining a marker for myofibrils (*UAS-Cherry-Gma*) with live mitochondria markers. Live dissection occasionally resulted in regions where myofibrils were mechanically severed (Fig. 3a). As shown above, areas with intact parallel myofibrils show elongated ellipsoid-shaped mitochondria with their long axis oriented in the direction of the myofibrils (Fig. 3b,d). Strikingly, severing myofibrils results in a dramatic rounding up of all neighbouring mitochondria into spheres (Fig. 3c,e). This transition was observed with both mitochondrial markers, MOM-GFP (Fig. 3c) as well as with mito-GFP (Fig. 3e). Interestingly, no obvious connection between the rounded mitochondria and the myofibrils remained visible within the severed area (Fig. 3e’’) strongly suggesting that mechanical pressure created by the tense myofibrils, rather than specific protein-protein binding, pushes mitochondria into their elongated shape covering the myofibrils.

**Fig. 3.**
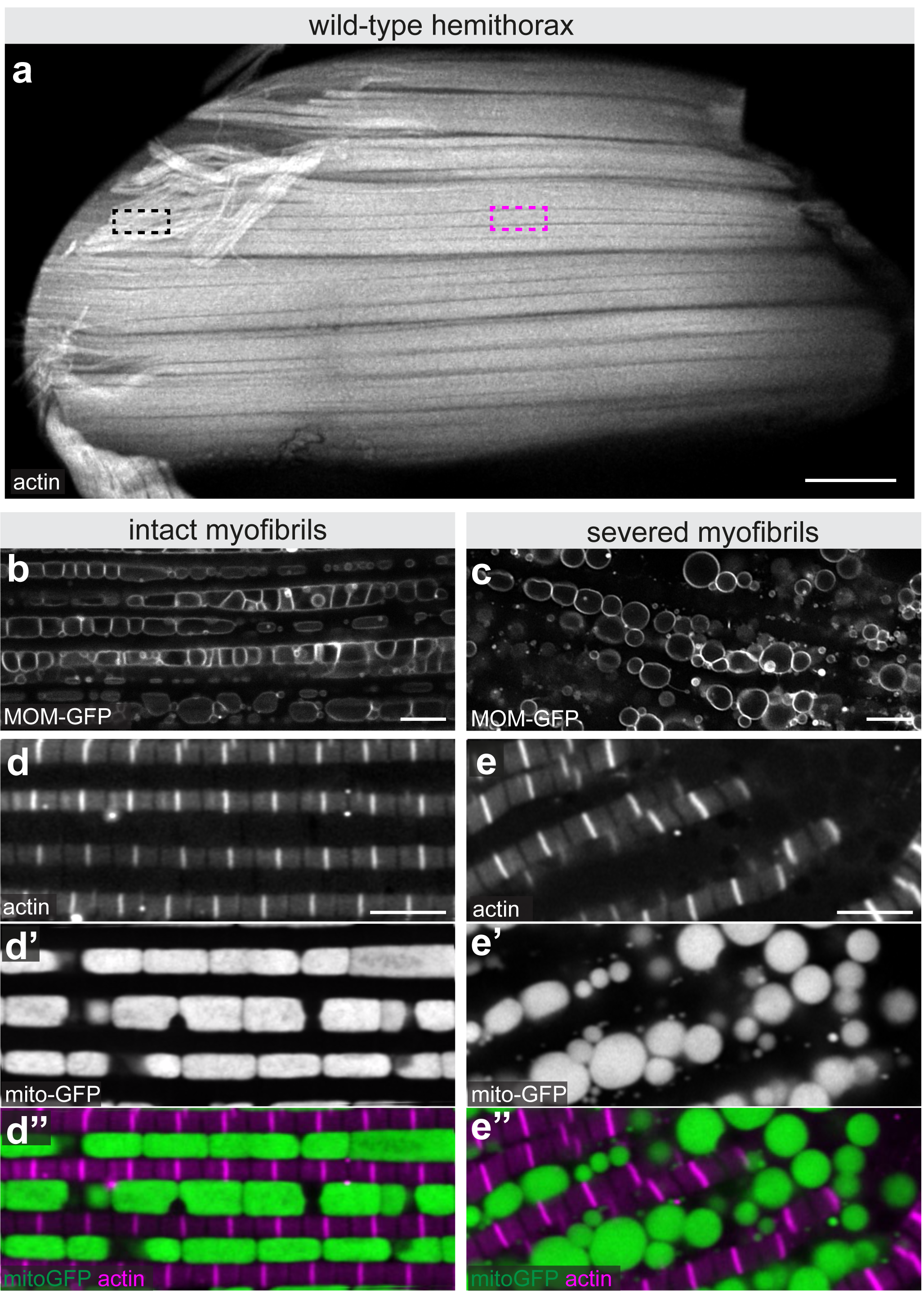
Myofibrils mechanically shape flight muscle mitochondria. (**a**) Live dissection of hemithorax in which actin has been labelled with Cherry-Gma expressed with *Mef2-*GAL4. Black rectangle indicates a severed area, in which myofibrils have been mechanically cut, magenta rectangle marks an intact area. (**b-e**) High magnification confocal sections of intact (b.d) and severed areas (c,e) of unfixed flight muscles in which mitochondria have been labelled with MOM-GFP (expressed with *Mef2*-GAL4) (b,c) or with mitoGFP together with Cherry-Gma to label myofibrils (d,e). Note the spherical mitochondria shape and their disengagement from the myofibrils in the severed areas. Scale bars are 100 µm (a) and 5 µm (b-e).

### Mitochondrial dynamics impacts myofibril development

Mechanical shaping of mitochondria by myofibrils should require a close contact between the two during muscle development. Mitochondria are highly dynamic organelles whose morphologies are defined by a delicate balance between mitochondrial fusion and fission^20-22^. Thus, we hypothesized that changing fusion or fission rates may not only change mitochondrial shapes but also impact on myofibril development. To test our hypothesis, we reduced mitochondrial fusion by knocking down *Mitochondrial associated regulatory factor* (*Marf*), a mitofusin required for outer mitochondrial membrane fusion in flight muscles^23,24^, with *Mef2*-GAL4 in muscles (*Mef2::Marf-IR*). Flight muscle fiber morphology of *Mef2::Marf-IR* flies is normal, however flight function, as assayed by a flight test, is impaired (Fig. 4a-c). As to be expected, reduced fusion rate results in smaller mitochondria in *Mef2::Marf-IR* flight muscles, which adopt a spherical instead of an ellipsoid shape (Fig. 4d,e,k). However, these small mitochondria intercalate normally between myofibrils resulting in wild-type shaped individualised fibrillar myofibrils with normal myofibril diameter and normal sarcomere length (Fig. 4d,e, Supplementary Fig. 3e,f). Also, the mitochondrial content is comparable to wild type (Fig. 4j). We observed the same phenotype when we increased mitochondrial fission rate during development by over-expressing *Dynamin related protein 1* (*Drp1*), a regulator of outer mitochondrial membrane fusion^23^ with *Mef2*-GAL4 (*Mef2::Drp1*) (Supplementary Fig. 3a-f). These data suggest that smaller mitochondria cannot sustain flight but are compatible with intercalating between myofibrils and thus enable normal fibrillar myofibril development of flight muscles.

**Fig. 4.**
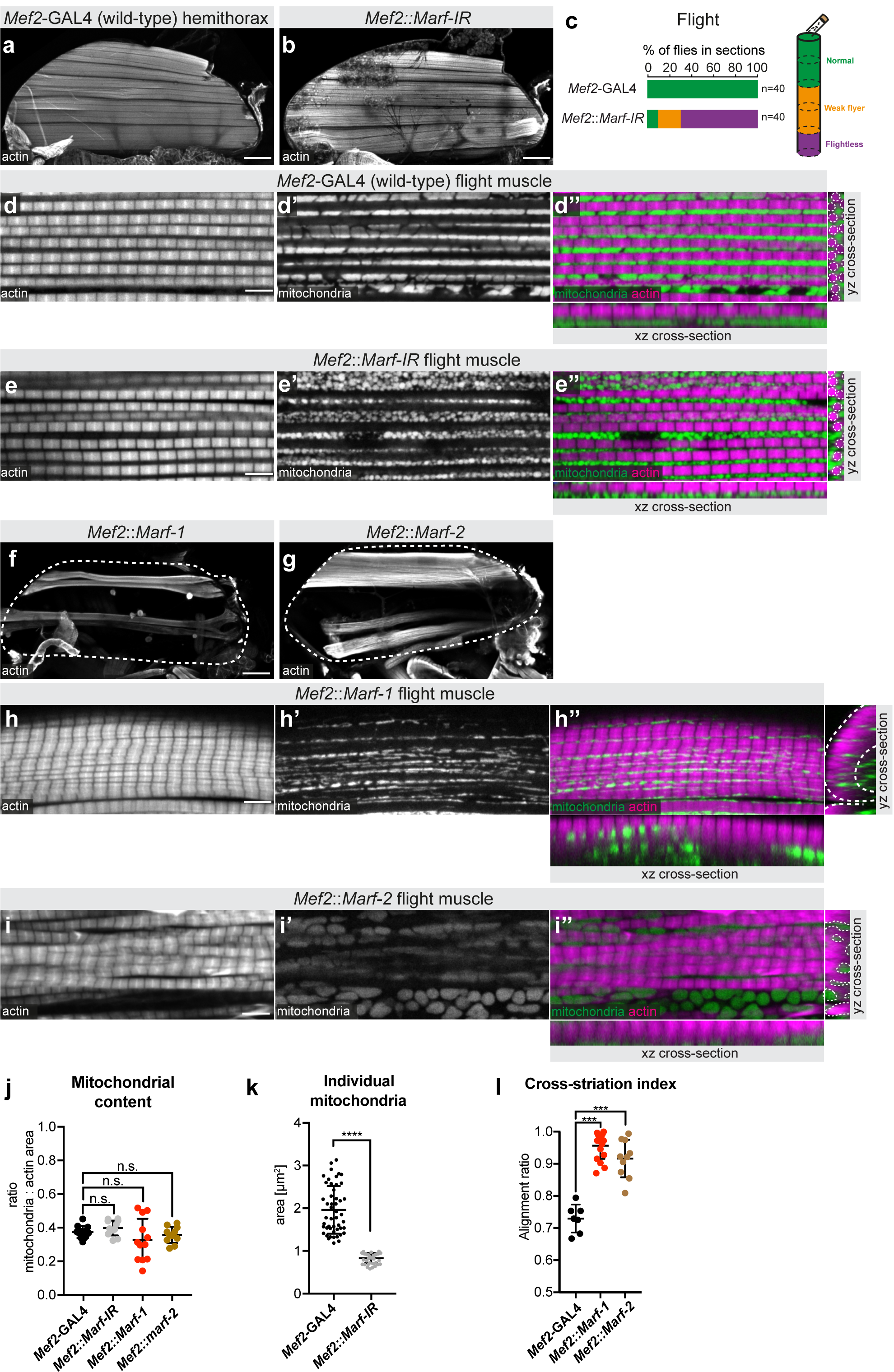
Mitochondria dynamics can impact myofibril development. (**a-i**) Adult hemithoraces (a,b,f,g), flight muscles (d,e,h,i) and flight test (c) of the indicated genotypes. Actin has been visualised with phalloidin, mitochondria with mito-GFP. White dashed lines outline the myofibrils in the yz cross-sections (d’’,e’’). Note the small round mitochondria present between normal myofibrils upon *Marf* knock-down (e). *Marf* overexpression using *UAS-Marf-1* or *UAS-Marf-2* causes fiber atrophy (f,g) and cross-striated myofibrils (h,i). Note that the mitochondria are largely separated from the aligned myofibrils, outlines by the dashed white line on the yz cross-sections (h’’,i’’). (**j-l**) Quantification of mitochondrial content (relative to actin area) (j), of individual mitochondrial area in a single confocal section (k) and the cross-striation index of the indicated genotypes (l). Significance from unpaired t-tests is denoted as p-values p ≤ 0,001 (***), p ≤ 0,0001 (****). (n.s.) non-significant. Scale bars are 100 µm (a,b,f,g) and 5 µm (d,e,h,i).

In an attempt to convert flight muscle mitochondria into larger networks, we performed the converse experiment and increased mitochondrial fusion by over-expressing *Marf* during development using two differently strong UAS-*Marf* constructs (*Mef2::Marf-1* and *Mef2::Marf-2*)^24,25^. In both cases, over-expression of *Marf* with *Mef2*-GAL4 results in fewer flight muscles, likely due to muscle atrophy during development (Fig. 4f,g). However, remaining flight muscle fibers show a dramatic change in their myofibril organisation with neighbouring myofibrils aligning laterally to cross-striated myofibrils, mimicking the cross-striated leg muscle morphology (Fig. 4h,i,l). The total mitochondrial content is similar to control flight muscles (Fig. 4j), however *Marf* over-expression results in an exclusion of mitochondria from the myofibril layer, similar to our observations in leg muscles (Fig. 4h’’,i’’). In some cases, in particular with the stronger *Marf-1* construct, perfect tubular muscles are generated with all myofibrils lining the outside of a tube and mitochondria located centrally (Supplementary Fig. 3g). This transformation from fibrillar to cross-striated myofibril morphology was also observed when mitochondria fission was suppressed by expression of dominant negative *Drp1* (*Mef2::Drp1k38a*)^26,27^ (Supplementary Fig. 3h-l) and thus is not a specific effect of *Marf* over-expression only but generally caused by tipping mitochondrial dynamics towards more fusion. Taken together, these results imply that increasing the mitochondrial fusion rate impacts myofibril development such that individual fibrillar myofibrils cannot form and instead fuse together to form cross-striated myofibrils.

### Mitochondrial fusion shifts transcription towards cross-striated fate

To decipher the mechanism of how a change in mitochondrial dynamics can impact myofibril morphology we investigated the expression of sarcomeric protein isoforms that are specifically expressed in fibrillar flight or cross-striated leg muscle types. We used GFP fusions of large genomic fosmid clones that recapitulate the endogenous expression dynamics with the added advantage to allow the quantification of expression levels without the need of antibodies^28^. Interestingly, we found that levels of both the flight muscle specific actin isoform Act88F-GFP and the flight muscle specific myosin binding protein Flightin (Fln-GFP) are strongly reduced in flight muscles overexpressing *Marf-1* (*Mef2::Marf-1*) (Fig. 5a-h). Conversely, Kettin (Kettin-GFP), a short isoform of the *Drosophila* titin homolog Sls, which is expressed at high levels in wild-type leg muscles^16^, is boosted in *Mef2::Marf-1* flight muscles (Fig. 5i-l). Hence, an increase in mitochondrial fusion during muscle development results in a transcriptional shift towards a more cross-striated muscle fiber type fate, which may contribute to the observed cross-striation phenotype.

**Fig. 5.**
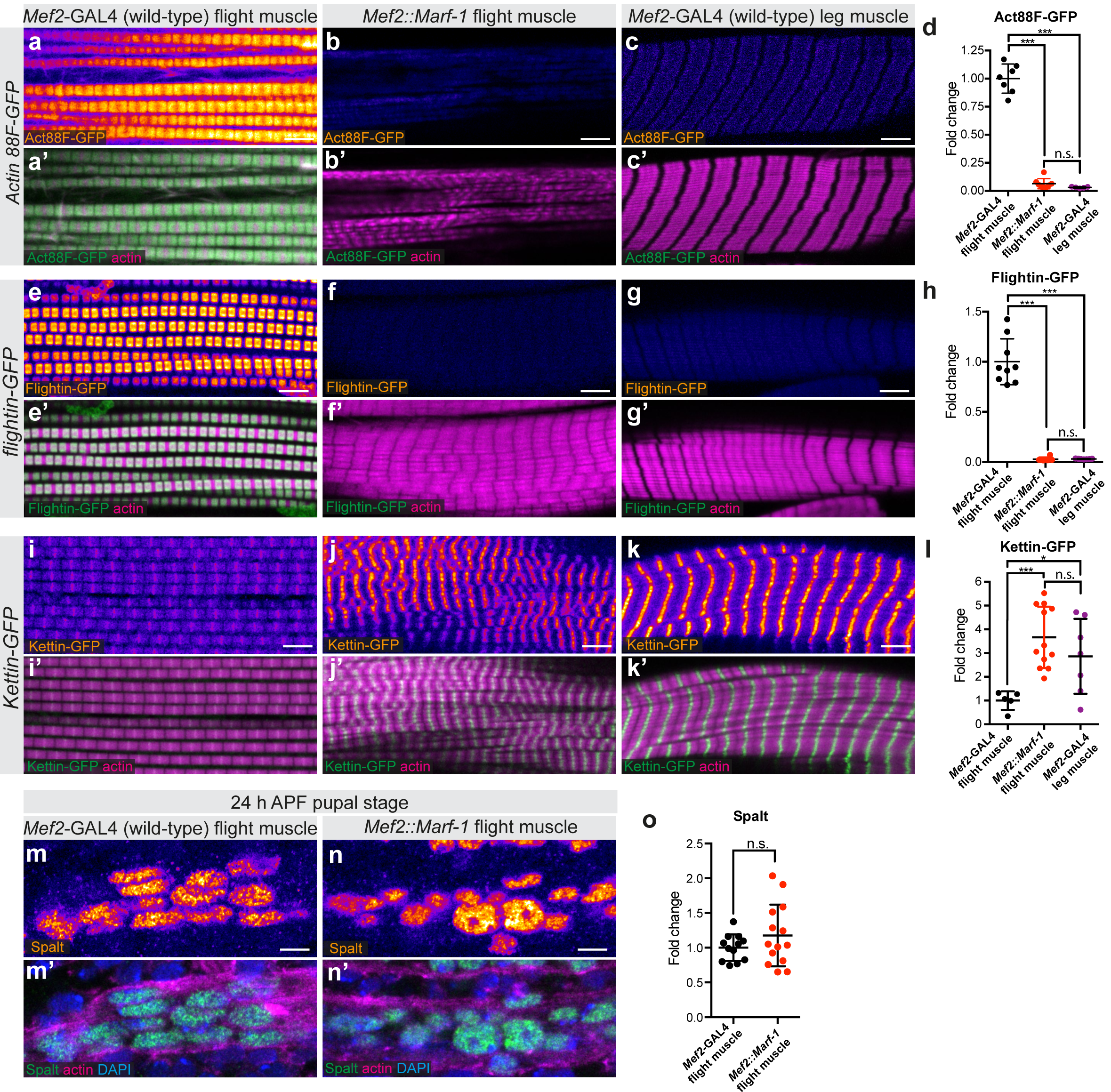
Mitochondria hyperfusion causes a transcriptional shift to cross-striated muscle type. (**a-k**) Adult wild-type (a,e,i) as well as *Mef2::Marf-1* flight muscles (b,f,j) and wild-type leg muscles (c,g,k) expressing GFP-tagged muscle-type specific proteins Actin 88F-GFP (a-c), Flightin-GFP (e-g) and Kettin-GFP (i-k); samples were fixed and actin was visualised with phalloidin. GFP fluorescence levels are represented via a heat map (fire LUT, white is higher). (**d**,**h**,**l**) GFP fluorescence was quantified with quantitative confocal microscopy (see Methods) and plotted relative to control flight muscle levels. Note that Marf over-expression in flight muscle converts the expression levels towards wild-type leg muscle levels. (**m-o**) Spalt protein levels in developing flight muscle myotubes at 24 h APF were quantified using immunostaining and quantitative confocal microscopy comparing wild type (m,o) to *Mef2::Marf-1*(n,o). Actin was visualised with phalloidin, nuclei with DAPI. Note the comparable expression levels. Significance from unpaired t-tests is denoted as p-values ≤ 0,05 (*), p ≤ 0,001 (***). (n.s.) non-significant. Scale bars are 5 µm.

A simple explanation of the observed phenotype would be that over-expression of *Marf* interferes with flight muscle fate patterning at an early stage of development. To investigate this, we quantified the expression levels of the Zn-finger transcriptional regulator Spalt, which was shown to be required and sufficient to induce fibrillar muscle fate^17^. Spalt is expressed at high levels immediately after myoblast fusion in the flight muscle myotubes^17^. Thus, we quantified Spalt protein expression during early flight muscle development at 24 h after puparium formation (APF) and found that Spalt levels in *Mef2::Marf-1* myotubes are comparable to wild type (Fig. 5m-o). This strongly suggests that flight muscle fate is induced normally in *Mef2::Marf-1* myotubes and, as consequence, that increased mitochondrial fusion impacts flight muscle development downstream of Spalt.

### Developmental timing of mitochondrial dynamics has differential impact on myofibril development

To better define the stage at which increased mitochondrial fusion can impact myofibril development, we tested a series of different GAL4-driver lines that are active at different stages of flight muscle development, in comparison to *Mef2*-GAL4 which is continuously active during all stages^29^. When *Marf-1* is over-expressed using *him*-GAL4 or *1151*-GAL4, restricting the over-expression to myoblasts and early stages of myoblast fusion, ending shortly after 24 h APF^18^, myofibrils and mitochondria show a wild-type morphology in adult flight muscle and flies can fly (Fig. 6a-e). This suggests that increasing mitochondrial fusion during myoblast and early myotube development, before myofibrils start to assemble, does neither impact myofibril development nor flight muscle function.

**Fig. 6.**
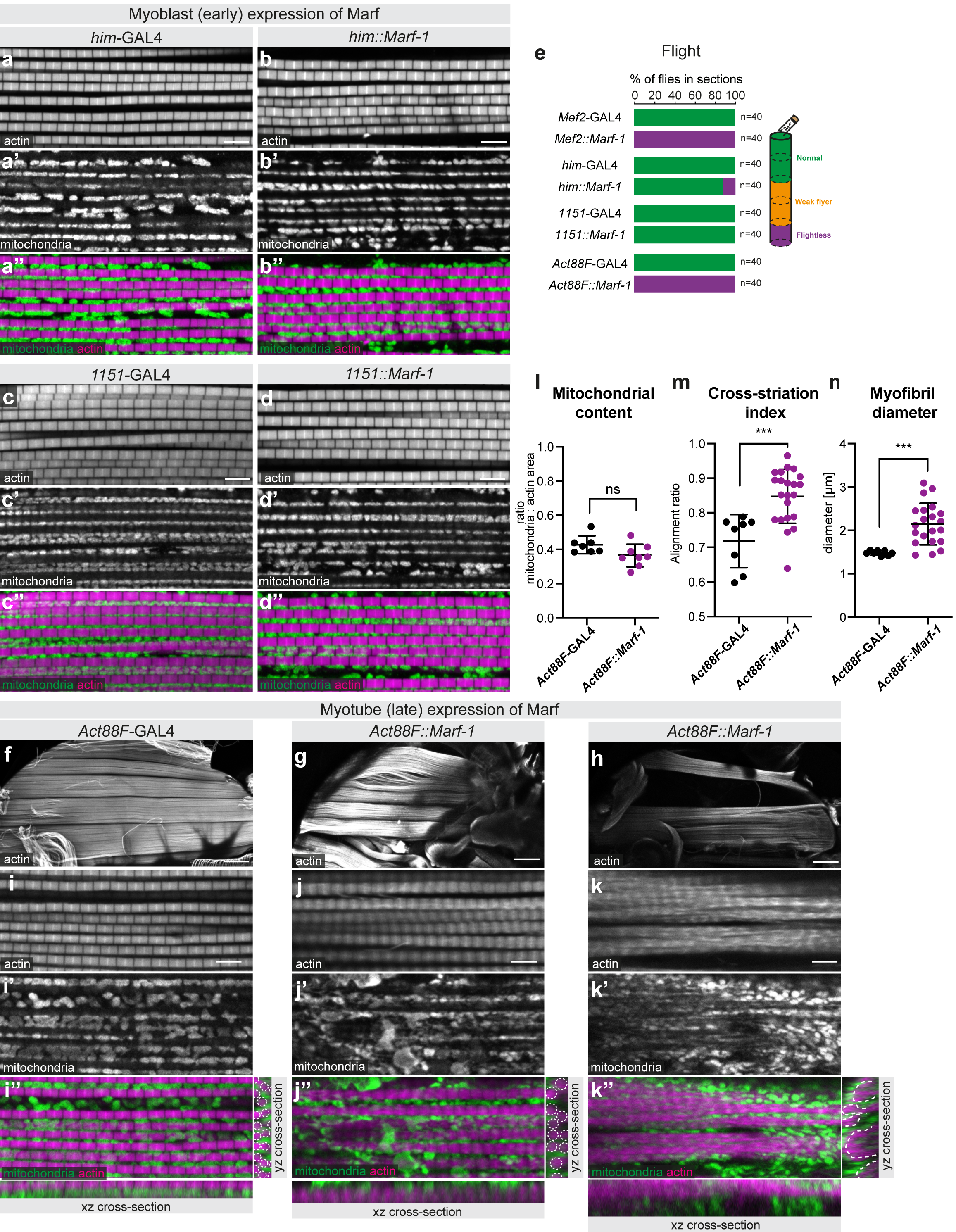
Developmental timing of mitochondrial dynamics impacts myofibril development. (**a-e**) Wild-type adult control (a,c) and early over-expression of *Marf-1* with *him*-GAL4 (b) or *1151*-GAL4 (d) flight muscles were stained with phalloidin and anti-complex V antibody to visualise myofibrils and mitochondria. Note the normal myofibril and mitochondria morphologies (b,d), which support flight (e). (**f-n**) Wild-type control (f) and *Act88F::Marf-1* hemithoraces (g,h), as well as flight muscles (i-k) expressing *Marf-1* during later developmental stages were stained with phalloidin and anti-complex V antibody to visualise myofibrils and mitochondria. Two representative phenotypes of *Act88F::Marf-1* flight muscles are shown, displaying either thicker myofibrils (indicated by white dashed outline in the cross-section) (j) or largely cross-striated myofibrils (k). Mitochondria are largely excluded from myofibrillar bundles in *Act88F::Marf* muscles (see dashed white outline of the myofibrils rich areas in k’’). (**l-n**) Quantification of the *Act88F::Marf* flight muscle phenotypes, quantifying mitochondrial content (l), cross-striation index (m) and myofibril diameter (n). Significance from unpaired t-tests is denoted as p-values ≤ 0,001 (***). (n.s.) non-significant. Scale bars are 5 µm (a-d,i-k) and 100 µm (f-h).

To over-express *Marf-1* during myofibril development we chose *Act88F*-GAL4, which is specifically and strongly expressed in flight muscles starting at about 24 h APF^30^ and thus includes all stages of myofibril assembly and myofibril maturation^31^. Strikingly, *Act88F::Marf-1* flight muscles show a severe myofibril phenotype, ranging from enlarged myofibril diameter (Fig. 6g,j,n) to partially cross-striated myofibrils (Fig. 6h,k,m). Although total mitochondrial content is similar to wild type (Fig. 6l) mitochondria are often excluded from the inter-myofibril space (Fig. 6j’’,k’’), thus creating space for myofibril diameter growth or myofibril alignment towards a cross-striated phenotype. As a consequence, *Act88F::Marf-1* flight muscles cannot support flight (Fig. 6e). We conclude from these results that induction of mitochondria hyperfusion specifically during stages of myofibril assembly and myofibril maturation strongly impacts myofibril growth and spacing.

### Mitochondria intercalate during myofibril assembly

We know little about the interplay between mitochondria and myofibrils during myofibril assembly and maturation stages. To explore this, we dissected wild-type flight muscles at 24 h APF, a stage at which a dense network of actin filaments is present, while myofibrils are still undefined. We found that mitochondria form a widespread filamentous network of tubules that is largely separated from the dense actin filament mesh at 24 h APF (Fig. 7a,b). When myofibrils have just assembled at 32 h APF, the mitochondria network has redistributed and mitochondria have intercalated between the myofibrils. As a consequence, myofibrils are individualised and are not in physical contact with neighbouring myofibrils (Fig. 7c,d). This finding is also supported by electron microscopy data that found mitochondria present between assembled myofibrils at 32 h APF^32^. This shows that mitochondria and myofibrils are present in close proximity directly after myofibrils have been assembled, suggesting a potential role of mitochondrial intercalation for fibrillar flight muscle morphogenesis.

**Fig. 7.**
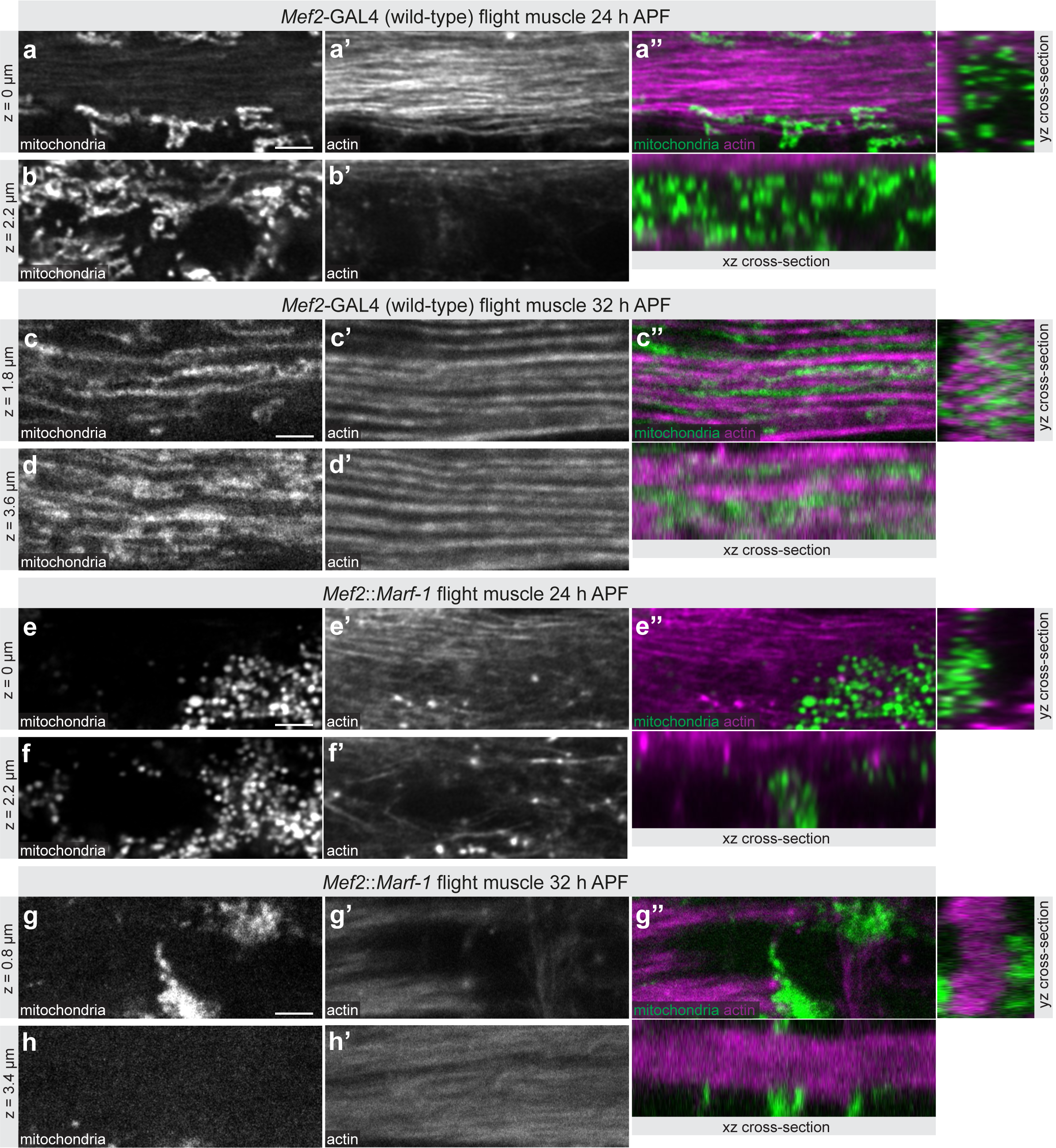
Developmental effect of mitochondria hyperfusion. (**a-h**) Developing wild-type flight muscles at 24 h after puparium formation (APF) (a,b) and 32 h APF (c,d), compared to *Mef2::Marf-1* flight muscles at 24 h APF (e,f) and 32 h APF (g,h). Mitochondria was visualised with mito-GFP and actin with phalloidin. Note the mitochondrial intercalation between myofibrils in wild-type 32 h APF flight muscles (c’’), which is blocked by *Mef2::Marf-1* (g’’). Scale bars are 2.5 µm.

### Fine-tuned mitochondria dynamics enables mitochondrial intercalation

To test the functional relevance of mitochondrial intercalation we explored myofibril and mitochondrial morphologies after over-expressing *Marf* with *Mef2*-GAL4. We found that continuous mitochondrial hyperfusion in flight muscles results in clustering of most mitochondria likely forming connected networks in a few areas outside of the actin filament mesh at 24 h APF (Fig. 7e,f). Interestingly, these mitochondrial networks are also maintained at 32 h APF preventing clustered mitochondria to intercalate between the forming myofibrils in *Mef2::Marf-1* flight muscles (Fig. 7g,h).

As myofibril development often appeared compromised when *Marf* was over-expressed with *Mef2*-GAL4, we wanted to explore the developmental phenotype in more detail using the late *Act88F*-GAL4 driver line. In *Act88F::Marf-1* flight muscles myofibrils assemble well at 32 h APF, but as in *Mef2::Marf-1*, most mitochondria clump together in large networks that are physically separated from the myofibril layer (Fig. 8a-d). Hence, mitochondrial intercalation is also strongly compromised in *Act88F::Marf-1* 32 h APF flight muscles.

**Fig. 8.**
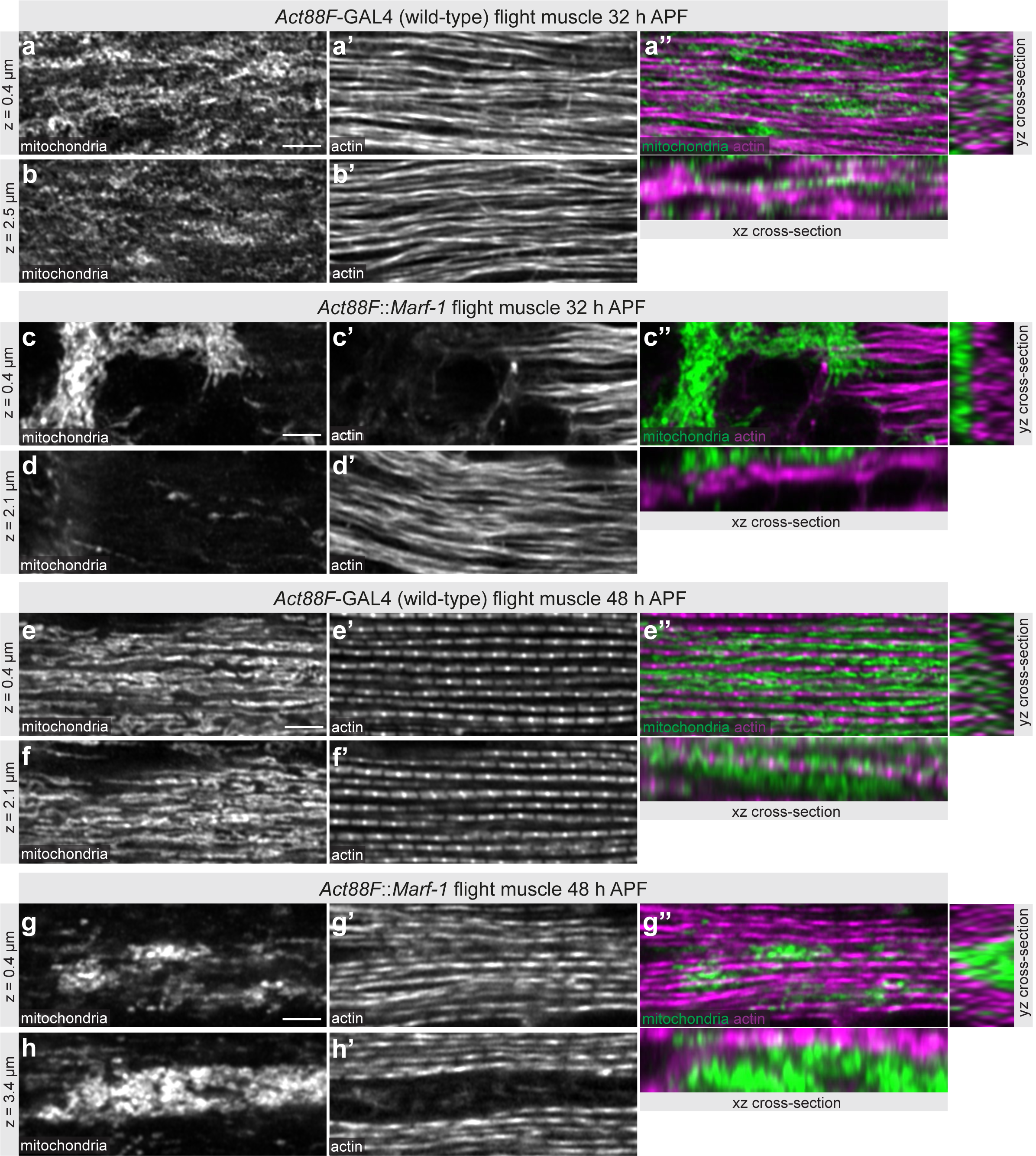
Mitochondria isolate individual myofibrils. (**a-h**) Developing wild-type flight muscles at 32 h after puparium formation (APF) (a,b) and 48 h APF (e,f), compared to *Act88F*::Marf-1 flight muscles at 32 h APF (c,d) and 48 h APF (g,h). Mitochondria was visualised by immunostaining against complex V (ATPase) and actin with phalloidin. Note how mitochondria isolate myofibrils in wild type (e’’) but fail to do so in *Act88F::Marf-1* with mitochondria clustering centrally (g’’). Scale bars are 2.5 µm.

Assembled myofibrils mature from 32 h to 48 h APF and display very regular sarcomeric patterns at 48 h APF^31^ (Fig. 8e,f). We wanted to investigate if the mitochondrial intercalation defect is maintained during myofibril maturation and how this impacts myofibril development. Indeed, we often found that mitochondria of 48 h APF *Act88F::Marf1* flight muscles stay networked in large clusters (Fig. 8g,h). These mitochondria clusters are sometimes even present in the center of a tube formed by closely aligned myofibrils (Fig. 8g’’). Thus, mitochondria physically separate the maturing myofibrils at 48 h APF in wild-type flight muscles, whereas hyperfused mitochondrial networks fail to do so. This provides a mechanistic explanation, why the intercalation block can result in myofibril diameter overgrow and often in lateral alignment of neighbouring myofibrils resulting in cross-striated fibers at the adult stage (see Fig. 9). Thus, balanced mitochondrial dynamics enables mitochondria to physically isolate the maturing myofibrils to support fibrillar flight muscle development.

**Fig. 9.**
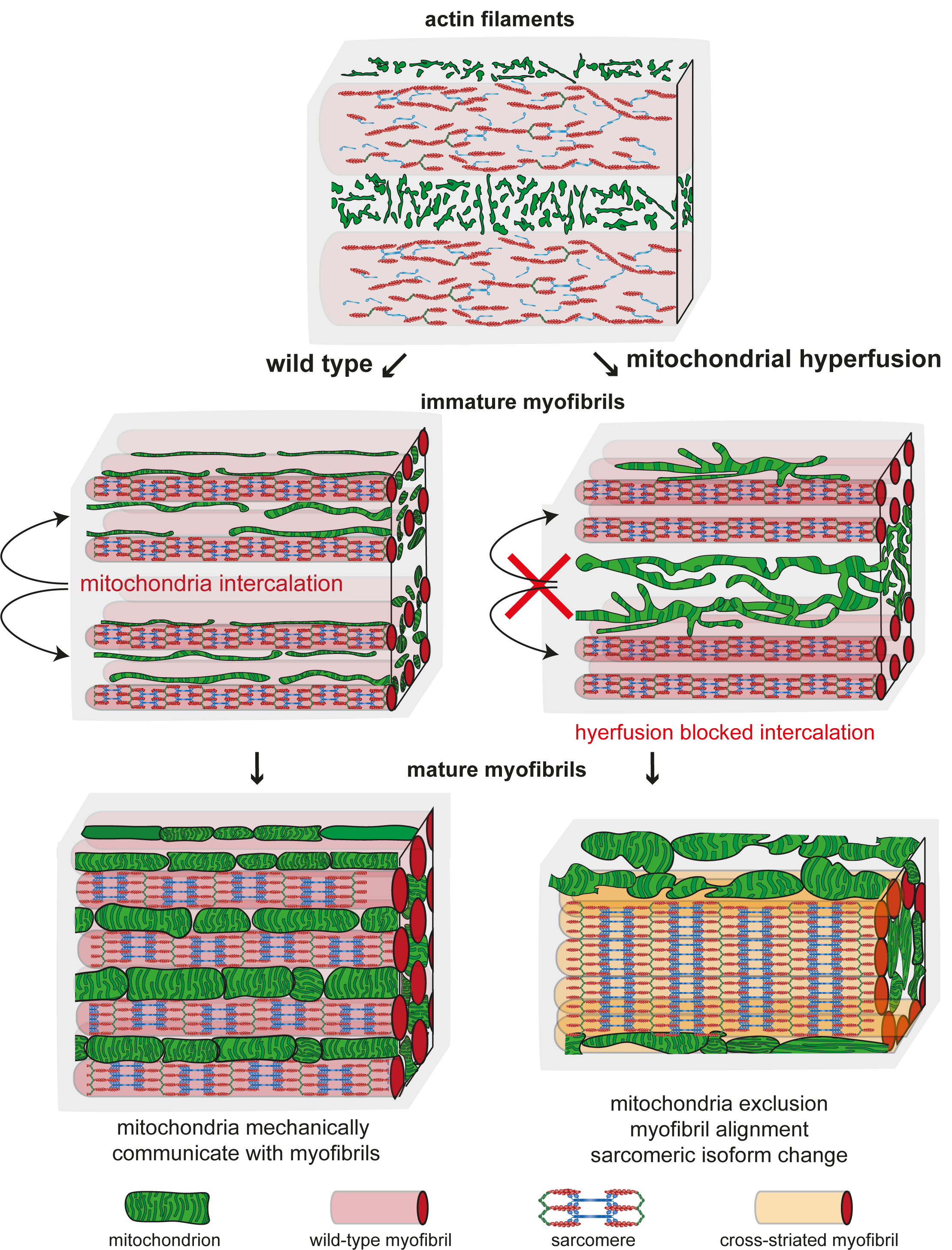
Mitochondria – myofibril communication model. Developing flight muscle schemes to highlight the interplay between mitochondria with actin filaments (top, early stage), immature myofibrils (middle, intermediate stage) and mature myofibrils (bottom, mature stage). Wild type is shown on the left and mitochondrial hyperfusion on the right. At the actin filament stage, mitochondria display a filamentous network morphology spatially separated from the actin filament mesh. Upon myofibril assembly, mitochondria intercalate between myofibrils in wild type and establish a tight mechanical communication. Myofibril and mitochondrial diameter growth causes generation of mechanical pressure and isolates individual myofibrils. Mechanical feedback ensures the correct myofibril diameter. In contrast, hyperfusion of mitochondria results in large interconnected mitochondrial networks that fail to intercalate between the immature myofibrils. As a consequence of mitochondrial exclusion, the mechanical communication to myofibrils is limited and myofibrils align with each other around centrally clustered mitochondria. Scheme for sarcomeric components was adapted from^3^.

## Discussion

Here we are proposing that mitochondria and myofibril morphogenesis are coordinated by a mechanical feedback mechanism in *Drosophila* flight muscles. The evidence for this hypothesis is five-fold. First, as soon as myofibrils have assembled they are surrounded by mitochondria, which isolate each of them from their neighbouring myofibrils. Hence, direct mechanical contact between neighbouring myofibrils is blocked (Fig. 9). Second, when myofibrils and mitochondria mature, both strongly expand in diameter, generating an extensive mechanical communication interface between them. The ellipsoid mitochondria shapes along the myofibril axis together with the induced mitochondrial indentations caused by the myofibrils strongly support a role for mechanical pressure from myofibrils on mitochondria and vice versa. Third, in contrast to leg muscles, no specific contact sites at particular sarcomeric locations are present in flight muscles arguing against localised protein-protein-interactions mediating the spatial proximity and hence favouring the mechanical interaction hypothesis. Fourth, relaxing mechanical constraints in the adult flight muscles by cutting myofibrils results in an immediate rounding of mitochondria, strongly suggesting that pressure directly shapes mitochondria. Finally, if intercalation is compromised, myofibrils grow larger in diameter, consistent with a mechanical feedback controlling myofibril diameter in flight muscles. Together, these data strongly support a role for mechanical forces coordinating mitochondria and myofibril morphogenesis in flight muscle (Fig. 9).

Surprisingly, we have found that interfering with mitochondrial intercalation changes the transcriptional state of the flight muscles by down-regulating some flight muscle specific sarcomeric isoforms and up-regulating at least one cross-striated muscle specific isoform (Fig. 9). Mechanistically, we showed that this change happens down-stream of Spalt, since Spalt-dependent flight muscle specification is normal. How does the defective mitochondria intercalation feedback on transcription? It has been shown that transcription is strongly regulated during myofibril maturation in flight muscles resulting in a boost of sarcomeric gene expression^31,33^. Furthermore, it is well established that during mammalian muscle fiber type maturation, sarcomeric isoform expression changes from embryonic isoforms, to neonatal ones and finally to adult isoforms^34,35^. How these switches are controlled is not fully understood, but it is conceivable that changes in mitochondrial metabolism may contribute. Alternatively, as manipulating mitochondrial dynamics affects myofibril alignment, this change in the biomechanical properties of the myofibrils may impact the transcriptional status of the muscle fiber. Together, such a transcriptional feedback would ensure a direct coordination between mechanical and physiological requirements of the developing muscle fibers.

How do mitochondria intercalate between myofibrils? This is likely an active mechanism as it happens rapidly during a few hours of muscle development. A first explanation could be that the driving force can either originate by the assembly of myofibrils directly, which re-distribute throughout the fiber, starting from a more peripheral actin filament meshwork^18,31^. A second more attractive explanation would be an active mitochondrial transport mechanism, as mitochondria align along the axis of the newly formed myofibrils. Transport could be achieved by microtubule motors, since they have been described to transport mitochondria in various other cell types, particularly in neurons^36,37^. Interestingly, microtubules have been found in close proximity to the freshly assembled myofibrils in flight muscles^32,38^ and hence ideally placed to mediate mitochondria intercalation and alignment with the myofibrils.

Mechanical roles of mitochondria are not limited to muscle fibers. Pushing forces of polymerising actin filaments against a mitochondria network surrounding the spindle in mouse oocytes demonstrated a mechanical role of mitochondria in spindle positioning^39^. Also in this system, a fine balance between mitochondrial fusion and fission was necessary for normal spindle positioning^39^. Similarly, mitochondrial remodelling into long giant mitochondria has been shown to be essential for sperm tail elongation during *Drosophila* spermatogenesis^40^. In these cells, mitochondria provide the platform for polymerising microtubules and the mechanical link between microtubules and mitochondria is essential for sperm tail elongation^40^.

Muscle fibers contain very crowded cellular environments. Thus particularly in cardiomyocytes, which do contain large amounts of mitochondria^9^ and share their high mechanical stiffness and high passive tension with flight muscles^41^, the mechanical communication between mitochondria and myofibrils might be most prominent. However, we found here that even in cross-striated *Drosophila* leg muscles mitochondria do contact sarcomeres similarly to the contacts described for the ‘intermyofibrillar’ mitochondria in proximity to the sarcomeric I-bands in mammalian muscle^7^. Thus, mechanical communication between myofibrils and mitochondria might be of general importance to successfully coordinate muscle development and homeostasis.

Interestingly, changing the fine-tuned fusion-fission balance of mitochondria results not only in severe muscle fiber phenotypes during mouse development^42,43^, but also leads to severe impairment of muscle function and fiber loss if acutely manipulated in adult mice^44,45^. Furthermore, maintaining a healthy balance of mitochondrial fission and fusion is also essential to build and maintain a healthy mouse heart^46^. Interestingly, reducing mitochondrial fission results in dilated cardiomyopathy in neonatal mouse hearts, coinciding with impaired myofibril morphogenesis^47^. Even manipulating mitochondria dynamics after birth causes cardiomyopathies in mice^48^. Together, this highlights the importance of mitochondrial dynamics for muscle development and maintenance. While it is recognised that mitochondria networks are highly dynamic in healthy and diseased muscle fibers and cardiomyocytes, the here hypothesized mechanical coupling between mitochondria networks and the contractile machinery is still underappreciated in mammalian muscle.

## Methods

### Fly strains and genetics

Flies stocks were maintained under normal culture conditions in humidified incubators with 12 hour light-dark cycles. All fly stocks were maintained on standard lab fly medium. The standard lab medium is a variation of the Caltech media recipe, which includes 8% (w/v) cornmeal, 2% (w/v) yeast, 3% (w/v) sucrose, 1,1% (w/v) agar, 1% (v/v) acid mix. To prepare the media, cornmeal (80 g), sucrose (30 g), dry-yeast (20 g) and agar (11 g) were mixed in 1 L of water and brought to boil with constant stirring. The media was allowed to cool down to 60°C, before 10 ml of acid mix was added. Acid mix was prepared by mixing equal volumes of 10% propionic acid (v/v) and 83.6% orthophosphoric acid. The medium was then poured in vials (∼10 ml/ vial) or bottles (50 ml/bottle) and allowed to cool down before storing at 4°C for later usage.

Wild-type control flies are either GAL4 driver crossed to *w[1118]* strain or the corresponding UAS lines crossed to *w[1118]*, as indicated in the figure legends. The strains used in this study are detailed in Supplementary Table 1. All crosses were developed at 21°C in order to reduce GAL4 activity, unless otherwise mentioned. Developmental times indicated are the equivalent to the characterised ones at 27 °C^31^.

### Generation of UAS-MOM-GFP transgenic flies

The *UAS-MOM-GFP* construct used in this study was generated by subcloning a gBlock (Integrated DNA Technologies) containing the *Drosophila* homologue of rat Tom20 minimal sequence determined as being sufficient for mitochondria outer membrane (MOM) targeting^49^(ATGATTGAAATGAACAAAACTGCAATCGGCATTGCAGCGGGAGTAGCTGGAACTCTGTTTATTGGATACTGCATCTACTTCGACAAGAAGCGCCGCAGCGATCCCGAGTACAAGAAGAAAGTCCGT), fused in frame to sfGFP into pUASt-attB vector^50^ with EcoRI and NotI. The resulting plasmid was sequenced with the following primers: 5’-[GCAGGCCGAATTCATGATTG]-3’ and 5’-[CGTGGTCAGCCATTAGAATG]-3’ and integrated into attP site VK00033 using standard methods^28^.

### Flight and leg muscle preparation

Preparations of adult hemithorax or fixed pupa for microscopy were performed as described^29^. Pupae were developed and staged at 21°C, and fixed by paraformaldehyde 4% in PBS with 0,3% Triton X-100 (PBS-T), for 25 minutes at 31 h, 42 h or 62,5 h after pupa formation (APF), which correspond to 24 h, 32 h or 48 h APF of development at 27°C, respectively^31^. Actin was labelled with phalloidin-rhodamine (1:500; Molecular Probes) and nuclei with DAPI. Mitochondria were labelled by the expression of GFP fused to a mitochondrial matrix (mito-GFP) or outer membrane signal (MOM-GFP) and detected by direct fluorescence without staining. Alternatively, mouse anti-complex-V (anti-ATP5a, Abcam ab14748; 1:500) immunostaining was used to label mitochondria. GFP-fosmid lines, as indicated in the fly strains table, were used for direct visualization of Kettin-GFP leg isoform, Flightin-GFP and Act88F-GFP protein levels^28^. Samples were washed twice with PBS-T (5min) and mounted in VectaShield containing DAPI using two cover slips as spacers.

### Flight

Flight tests were performed as previously described^51^. Twenty males one week old were collected at least 24 h prior to testing and then dropped into a 1 m long plexiglass tube and with 8 cm inner diameter, divided into five zones. Those that landed in the top two zones were considered ‘normal fliers’, those in the next two zones ‘weak fliers’ and those that fell to the bottom of the cylinder ‘flightless’. In crosses with GAL4 insertions on the X chromosome females were used. Tests were repeated at least twice per genotype, for a minimum of 40 flies in total per condition.

### Live dissection to visualise flight muscles without fixation

Living hemithoraces were dissected and mounted in Schneider medium. Living samples were imaged within 30 minutes following dissection. Dissection consisted of a precise incision through the cuticle with sharp forceps (#11252-20 Dumont#5, Fine Science Tools) at the median plane resulting in the separation of the two hemi-thoraces. Ventral connective tissues were cut along the midline into two halves using fine dissection scissors (#15009-08 Fine Science Tools) to completely detach left from right halves. The dissection is usually non invasive for the flight muscle resulting in intact flight muscle fibers attached to the tendon cells of the thorax. Samples were mounted in Schneider medium using two cover slips as spacers and imaged immediately.

### Cross-striation Index

To quantify the vertical alignment of individual myofibrils we a defined a “cross-striation index” as the ratio between the distance needed to connect M-bands from neighbouring myofibrils and a straight line perpendicular to the myofibril horizontal axis from first to last myofibril used for quantification. To avoid bias, the nearest M-band was chosen when a horizontal path needs to be made in between myofibrils. Perfect alignment results in a ratio of 1 and lower values represent progressively less alignment (see Supplementary Fig. 1).

### Sarcomere quantification

Sarcomere length and diameter quantification were made using the MyofibrilJ plugin for Fiji (https://imagej.net/MyofibrilJ)^31,52^. For genetic interventions that result in strong sarcomere phenotypes, which cannot be analysed automatically by the plugin, measurements were made manually. An average of ten myofibril diameters per sample was then made on a interpolated YZ projection using Fiji. If samples showed a strong cross-striation phenotype, they were not quantified for myofibril diameter, but instead included for the cross-striation index quantification.

### Mitochondria content quantification

Total areas of actin (phalloidin) or mitochondria (mito-GFP) were identified via Otsu thresholding in Fiji for each individual acquisition channel. This was done for each Z-plane and channel, and thresholding was reset for each new plane in the same image, to correct for signal loss due to section depth. Multiple quantifications (the entire Z-stack at multiple XY ROIs per fly) from single flies were averaged and plotted as a single value in the figure plots (each fly counts as n=1).

For tubular leg muscles and *Marf* gain of function flight muscles, an interpolation across the Y-axis was made to generate a new image in which the Z-axis becomes the longitudinal axis of the tube (Z-depth cross-section). This new stack was then segmented for each slice (1024 in total for each image stack) to distinguish the peripheral area of the tube - rich in actin -, from the central area where a higher amount of mitochondria and no actin signal are present. Signal from either channel, actin or mitochondria, was then quantified as described above.

### Individual mitochondrion area

Mitochondria signal from mito-GFP expressing adult flight muscles was used to individually segment each mitochondrion across all Z-planes from the entire Z-stack. Fiji was used to apply a Gaussian filter (lambda=2), background correction, and Otsu thresholding, followed by watershed on the binary image. Individual objects were quantified for area (6,894 for wild type and 14,443 for *Mef2::Marf-IR* in total).

### 3D reconstruction and analysis

High-resolution confocal imaging was performed using a Zeiss LSM880 confocal microscope equipped with an Airyscan detector. Mitochondria were visualised in flight and leg muscle with two different labels: 1) mitochondria matrix mito-GFP or 2) mitochondrial outer membrane (MOM)-GFP. Flight muscle mitochondria were then segmented with a machine learning algorithm described in detail below, leg muscle mitochondria were segmented with the Fiji plugin “Interactive Watershed” (https://imagej.net/Interactive_Watershed). This plugin allows for extensive manual optimization of objecting splitting/joining in large stacks and includes 3D water-shedding, essential for our 3D reconstruction. Continuous validation for the watershed splitting was verified manually and we opted to have more splitting than to have too many large objects (by missing splitting). This compromise led to some network connections between mitochondria being missed.

The resulting binary images, for both tissues, were then connected in 3D via the MorpholibJ plugin for Fiji using ‘connected components labelling’^53^. Size Open (min 100 voxel filter) was applied and objects on the borders of the 3D space were discarded, and a 26 voxel connectivity used between Z-slices. This was followed by 3D object analysis in MorpholibJ, from which we obtained individual volumes used to colour code the mitochondria in the volume renderings, as well as the ellipsoids, long, medium and short axis and azimuts. 3D visualisation was done with the 3D Viewer plugin from Fiji and with Amira Software (Fisher Scientific).

### Deep Learning segmentation

A Deep Learning model has been trained to segment mitochondria labelled with MOM-GFP. A fully convolutional encoder-decoder UNET architecture has been used with an ImageNet pre-trained seResNet18 encoder as backbone^54,55^. To train the model, we built a relatively small dataset consisting of only 13 pairs of 128×128 image tiles extracted from one part of the entire 1024×1024 image stack and their corresponding manually drawn masks with Fiji. To perform strong network regularization in order to increase model performances despite the small dataset size, a data augmentation approach was successfully applied to virtually increase the training set size by generating, for every epochs (20 epochs in total), 400 batches of 32 artificially generated 128×128 image tiles by using a combination of random horizontal/vertical flips, width/heights shift and zooms of the original training dataset.

Dice-Sörensen loss function were chosen to optimize the network weights with Adam optimizer (learning rate 1e-4) and Intersection Over Union metrics has been used to assess segmentation quality (0,85 IoU on validation set). The model has been trained using Python Keras framework (version 2.2.4) with Tensorflow (version 1.15) as backend on one Nvidia Quadro GV100 GPU card.

Each slice was segmented individually by splitting every slice into 64 adjacent tiles of size 128×128 to feed the trained model, retrieve their predicted segmentation mask and recombine everything to achieve the whole slice and volume segmentation. The resulting binary images were then connected in 3D via the MorpholibJ plugin for Fiji as described above. (See Supplementary Fig. 2a,b).

### 3D object classification

Classification of objects as spheres, ellipsoids or rods was performed according to ^56^. Elongation descriptor was calculated as the length of object’s ellipsoid length divided by half of the sum of its width and thickness. Flatness descriptor was obtained by dividing object’s width by its thickness. Sphere class was associated to every object having both their elongation and flatness less than 1.3; rod class was assigned to object having an elongation superior to 2.5; and all remaining objects were assigned to the ellipsoid category. The classes were used to colour code individual mitochondria and we verified for their accuracy by going through the stacks manually.

### Quantifications and Statistical Analysis

Detailed information on the number of animals/samples used for each quantification shown in the figures, as well as the statistical tests and p-values, is presented in Supplementary Table 2.

### Serial block face electronic microscopy

One-week-old *Drosophila* thoraxes were dissected rapidly in cold PBS and immediately fixed in 2% paraformaldehyde, 2.5% glutaraldehyde over night (ON) at 4°C. Samples were contrasted for 1 hour in 3% osmium, 20 minutes in TCH, then 30 minutes in osmium before being incubating at 4°C ON in 1% uranyl acetate. Between each incubation, five times 3-minute washes in water were done. The samples were incubated for 30 minutes in fresh aspartate and then dehydrated by a series of one-hour consecutive incubations in 20%, 50%, 70%, 90%, 100% acetone. Then, the samples follow a series of incubations of 2 h room temparature in a resin (Durcupan 25%, 50%, 75% and 100% diluted in acetone). The Durcupan 100% is renewed and incubated for 16 hours, then incubated for 48 hours at 60°C for the polymerization of the resin. Details are described in the NCMIR protocol for SBF-SEM^57^. Imaging was carried out on an FEI Teneo VS running in low vacuum (50 Pa), at 2kV and using a backscattered electrons detector. The acquisition voxel size was 5×5×40 nm. The segmentation was carried out manually in IMOD.

## Acknowledgements

The authors are indebted to Nicolas Broully and the IBDM imaging facility for help with image acquisition, maintenance of the microscopes, 3D visualisation and generating animations and the Turing Centre for Living Systems engineering team for their help with data analysis. The electron microscopy experiments were performed on the PiCSL-FBI core facilty (Nicolas Broully), we acknowledge the France-BioImaging infrastructure supported by the French National Research Agency (ANR– 10–INBS-04-01, Investments for the future). The authors are grateful to Christophe Pitaval and Céline Guichard for fly embryo injections and to Flybase for database organisation. Fly stocks obtained from the Bloomington *Drosophila* Stock Center (NIH P40OD018537) and the Vienna *Drosophila* Resource Center (VDRC) were used in this study. The authors thank Bianca Habermann, Pierre Mangeol, Qiyan Mao and Clara Sidor for helpful discussions and critical comments for this manuscript.

## Author contributions

J.A., N.M.L. and F.S. conceptualised and designed the study; J.A. performed most of the experimental work and image acquisition with help by C.R. for the developmental data; J.A. and T.R. generated the serial-block face electronic microscopy data; F.D. and N.M.L. performed the 3D reconstruction analysis; F.D. developed and implemented the machine learning segmentation analysis; J.A., N.M.L. and F.S. analysed the data and prepared the figures; N.M.L. and F.S. wrote the manuscript with input from all authors.

