## Supplementary Figures and Legends for "Myofibril and mitochondria morphogenesis are coordinated by a mechanical feedback mechanism in muscle"

**Quantification of the cross-striation index**

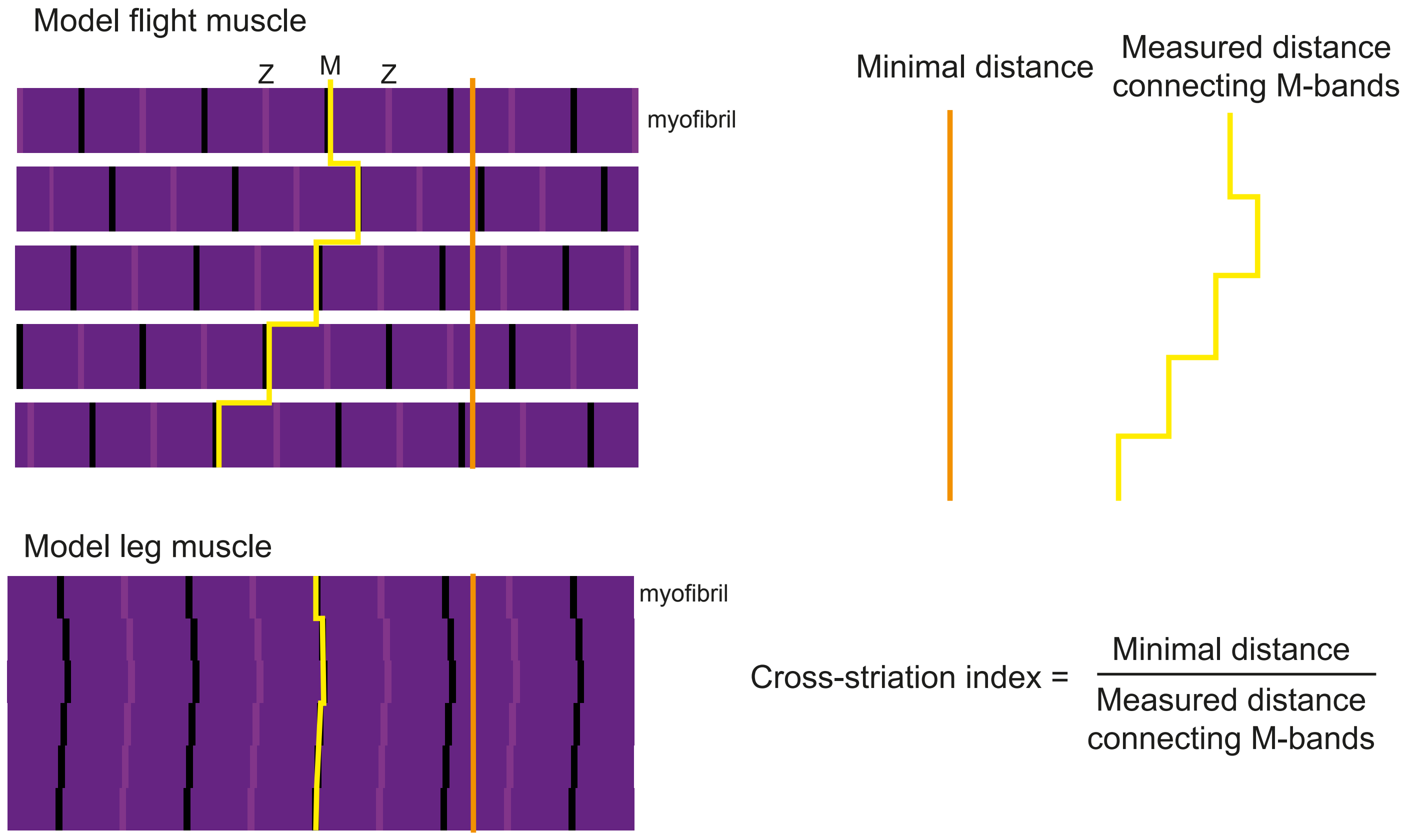


**Supplementary Figure 1**

**Supplementary figure legends.**

**Supplementary Fig. 1 Cross-striation index.** A cross striation index was defined as the ratio between the path length to connect M-bands (M) from adjacent myofibrils (yellow path) and the corresponding length of a straight line perpendicular to the myofibril axis (orange path). The closest M-band was chosen for the path connecting two myofibrils. Perfect alignment results in a cross-striation index of 1, lower values represent progressively weaker alignment.

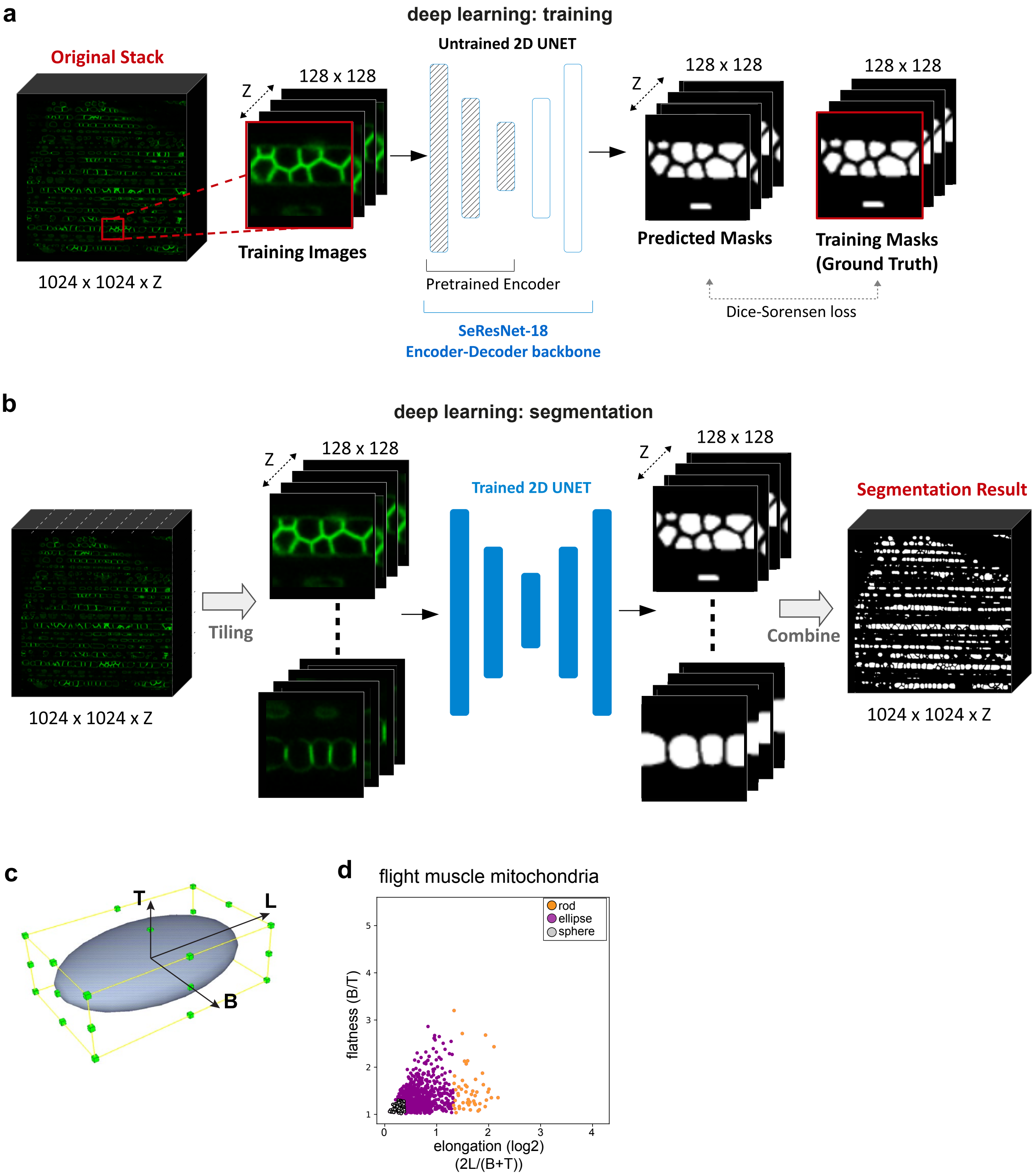


**Supplementary Figure 2**

**Supplementary Fig. 2 Mitochondrial shape annotation. (a)** Deep learning training: One small patch with 128x128 pixel resolution was extracted from one confocal stack (MOM-GFP labelled mitochondria) and was manually segmented using Fiji to generate a set of images and their corresponding ground truth masks. Training has been achieved through a 2D UNET network architecture based on the SeResNet18 backbone with an encoder part pretrained on ImageNet Database using a Dice-Sorensen loss to optimize the network weights and Intersection Over Union (IoU) metrics to assess the segmentation quality. **(b)** Deep learning segmentation: four confocal stacks (MOM-GFP labelled mitochondria) with 1024x1024 pixel resolution were processed separately. For each of them, the whole stack has been has been divided into 64 small 128x128 patches (tiling) and processed individually by the trained SeResNet18 2D UNET to generate a set of segmented patches which have further been combined to build the whole 1024x1024 stack segmentation result. (**c**) Diagram illustrating the long "L", middle "B" and short "T" axis of a model mitochondrium used for shape classification. The angle between the long axis relative to the myofibril axis was used to assign mitochondria orientation. (**d**) Morphology comparison on the basis of morphological descriptors, elongation and flatness of flight muscle; each dot represents an individual mitochondrium from a single 3D reconstruction (total volume of 30,300 $\mu m^3$).

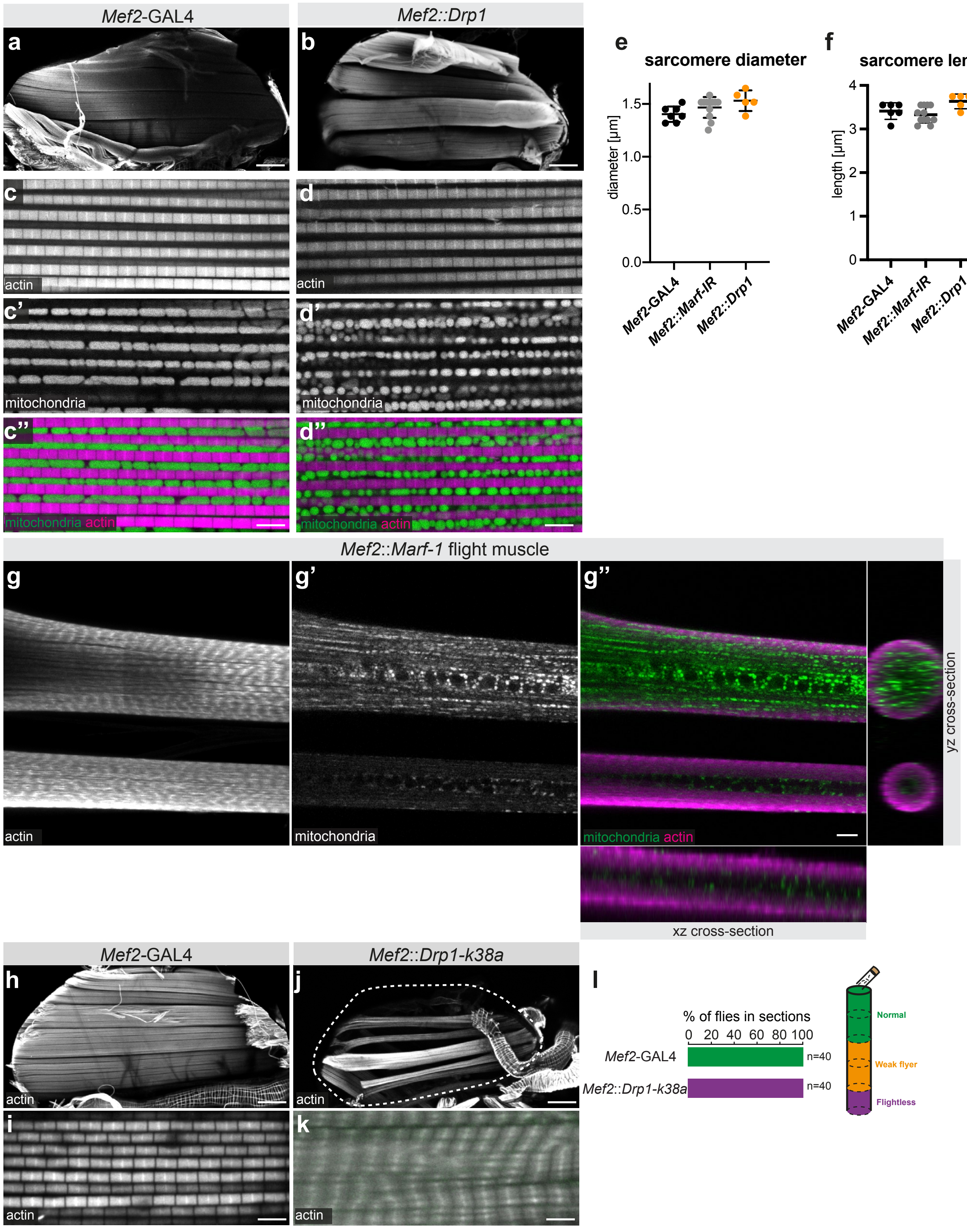


**Supplementary Figure 3**

**Supplementary Fig. 3 Manipulation of mitochondrial dynamics.** (**a-d**) Adult hemithoraces (a,b) and flight muscles (c,d) from *Mef2*-GAL4 (a,c) and *Mef2::Drp1* (c,d), in which actin has been visualised with phalloidin and mitochondria with mito-GFP. Note the smaller round mitochondria upon *Drp1* over-expression (d). (**e,f**) Quantification of myofibril diameter (e) and sarcomere length (f) of the indicated genotypes. (**g**) Thick confocal stack displaying two *Mef2::Marf-1* flight muscles displaying a cross-striated tubular morphology. Note the centrally located mitochondria. (**h-l**) Adult hemithoraces (h,j) and flight muscles (i,k) from *Mef2*-GAL4 (h,i) and *Mef2::Drp1-k38a* (j,k), in which actin has been visualised with phalloidin and mitochondria with mito-GFP. Note the aligned myofibrils (k) and the flightless phenotype upon expression of *Drp1-k38a* (l). Scale bars are 100 µm (a,b,h,j), 10 µm (g) or 5 µm (c,d,i,k).

**Supplementary Table 1** List of all fly strains and reagents used in this study.

**Supplementary Table 2** Data providing all number of animals and samples used for all the quantifications plotted in the figures. Where possible, individual values for the quantifications are presented, or averages per animal/sampled area, for each "n". Statistical analysis and p-value calculations are included.

**Supplementary Movie 1 (associated with Fig. 2):** Animation from the rendering shown in Fig. 2c of flight muscle mitochondria, distributed along the longitudinal axis of the myofibrils. Individual mitochondria are coloured randomly to highlight their spatial distribution .

**Supplementary Movie 2 (associated with Fig. 2):** Animation from the rendering in Fig. 2i,j showing the 3D reconstruction using images acquired each 40 nm via serial block-face electron microscopy. Individual mitochondria are shown with random colour and myofibrils appear during the movies in magenta. A single mitochondrium is highlighted in light pink together with myofibrils at the end of the movie.

**Supplementary Movie 3 (associated with Fig. 2):** Animation from the rendering show in Fig. 2k. Leg muscle mitochondria are labelled with mito-GFP.

**Supplementary Movie 4 (associated with Fig. 2):** Animation from the rendering shown in Fig. 2m showing segmented leg muscle mitochondria. Separated mitochondria are coloured randomly.

**Supplementary Movie 5 (associated with Fig. 2):** Animation from the rendering shown in Fig. 2n showing a close up of the leg muscle mitochondria network. Note the extensions that protrude from mitochondria into the space above and below where myofibrils are located (not represented).

**Supplementary Movie 6 (associated with Fig. 2):** Animation from the rendering shown in Fig. 2o displaying the complex shape of a single leg muscle mitochondrium
